# The human secretome – the proteins secreted from human cells

**DOI:** 10.1101/465815

**Authors:** Mathias Uhlen, Hanna Tegel, Åsa Sivertsson, Chih-Chung Kuo, Jahir M. Gutierrez, Nathan E. Lewis, Björn Forsström, Melanie Dannemeyer, Linn Fagerberg, Magdalena Malm, Helian Vunk, Fredrik Edfors, Andreas Hober, Evelina Sjöstedt, David Kotol, Jan Mulder, Adil Mardinoglu, Jochen M. Schwenk, Peter Nilsson, Martin Zwahlen, Jenny Ottosson Takanen, Kalle von Feilitzen, Charlotte Stadler, Cecilia Lindskog, Fredrik Ponten, Jens Nielsen, Bernhard O. Palsson, Anna-Luisa Volk, Magnus Lundqvist, Anna Berling, Anne-Sophie Svensson, Sara Kanje, Henric Enstedt, Delaram Afshari, Siri Ekblad, Julia Scheffel, Borbala Katona, Jimmy Vuu, Emil Lindström, LanLan Xu, Roxana Mihai, Lucas Bremer, Malin Westin, Muna Muse, Lorenz M. Mayr, Sinead Knight, Sven Göpel, Rick Davies, Paul Varley, Diane Hatton, Ray Fields, Bjørn G. Voldborg, Johan Rockberg, Lovisa Holmberg Schiavone, Sophia Hober

## Abstract

The proteins secreted by human tissues (the secretome) are important for the basic understanding of human biology, but also for identification of potential targets for future diagnosis and therapy. Here, we present an annotation of all predicted secreted proteins (n=2,623) with information about their spatial distribution in the human body. A high-throughput mammalian cell factory was established to create a resource of recombinant full-length proteins. This resource was used for phenotypic assays involving β-cell dedifferentiation and for development of targeted proteomics assays. A comparison between host cells, including omics analysis, shows that many of the proteins that failed to be generated in CHO cells could be rescued in human HEK293 cells. In conclusion, the human secretome has been mapped and characterized to facilitate further exploration of the human secretome.

---

We have initiated a comprehensive study of proteins secreted by human cells with the objective to map and classify them, as well as to generate a physical resource based on production in cultured mammalian cells and to use this resource of purified recombinant proteins for various studies, including phenotypic screens. Here, the human secretome is defined according to the Human Protein Atlas classification^1,2^ with proteins having a signal peptide, but lacking a transmembrane region. According to these criteria, a list of approximately 2600 potentially secreted protein in humans was assembled. This list of potentially secreted proteins in humans is a useful resource for all researchers interested in secreted proteins as targets for diagnostic and therapeutic drugs. Here, we report on a new program denoted the “Human Secretome Project” in which synthetic constructs for each of these genes have been generated and used for protein expression in mammalian cell factories.

A particularly interesting subgroup of the human secretome is the proteins that translocate to peripheral blood, thus having systemic effects in the human body. Blood proteins are attractive to study, since they are involved in many important body functions, such as inflammation, coagulation, defense, signaling and blood homeostasis, and they are also of great interest as possible molecular targets for clinical diagnostics and precision medicine. Importantly, although the presence of a signal peptide directs protein translocation across the Endoplasmic Reticulum (ER) membrane in conjunction with protein synthesis, not all of these proteins end up in the blood. The final fate of the protein depends on many factors, including retention signals, such as the 4 amino acid long C-terminal ER-retention signal KDEL^3^, the 3 or 9 amino acid long peroxisome targeting signal^4, 5^ and the 10-70 amino acid long mitochondrial target signal sequences^6^. Proteins with retention signals thus end up intracellularly despite the fact that they belong to the human secretome. In addition, other proteins are secreted to the gastrointestinal tract, such as pancreatic enzymes and proteins secreted from the salivary glands. Furthermore, many proteins are secreted to the proximity of the cell of origin. This includes matrix proteins and many proteins expressed in gender specific tissues and specialized tissues, such as the eye. Here, the number of proteins in each of these groups have been predicted to allow stratification of the human secretome with regards to the final localization of the protein.

A knowledge resource has been created, including both classification with regards to the predicted spatial location in the human body, and body-wide transcriptomics data showing the cellular and/or tissue origin of each of the secretome proteins. In addition, expression data is presented for all the gene constructs in the CHO cell factory to allow in-depth analysis of the relationship between protein sequence and yield. For some representative CHO expression clones, including both high and low producers, omics data have been generated and data is also presented using a human cell line HEK293 for a selection of the clones with low or no production in CHO. In conclusion, we will provide an open access knowledge resource to facilitate basic and applied research covering the proteins actively secreted in human cells, tissues and organs.

## Results

### Defining the human secretome

An analysis according to Uhlen et al^1^ was performed to identify all protein-coding genes with a signal peptide and no transmembrane region. Also proteins annotated as secreted by UniProt (www.uniprot.org), were added to this list. We excluded the complex immunoglobulin genes and proteins with no corresponding entry in the current Ensembl version (v92). The number of genes encoding potentially secreted proteins (the human secretome) were found to be 2,623 which is approximately 13% of all human protein-coding genes. Based on literature, bioinformatics and experimental evidence, these protein-coding genes were subsequently annotated individually for their spatial distribution in the human body (**Fig. 1A**). 625 proteins were identified as blood proteins, with a fraction of these having other main location, 515 were annotated as secreted to local compartments, e.g. male/female tissues, brain or other local tissues such as the eye or the skin. 76 proteins were identified as secreted to the gastrointestinal tract, including 26 produced by the salivary glands. Many proteins are also involved in the forming and function of the extracellular matrix, including laminins, collagens, elastin and fibronectin and more than 200 proteins were annotated to this class of secreted proteins. Approximately thousand proteins were annotated to be intracellular/membrane proteins, although it is not unlikely that some of these have isoforms actively secreted to blood, thus calling for more in-depth analysis of these proteins. Finally, 193 genes were coding for proteins lacking supporting data for their location. These are interesting proteins for studies to explore their function and location. In summary, the annotation has classified the human secretome into various groups based on their final location and the analysis suggests that the blood proteins consisting of proteins actively secreted to the blood (n=625) constitute a small fraction (24%) of the human secretome and only 3% of the total protein-coding genes in humans.

**Figure 1.**
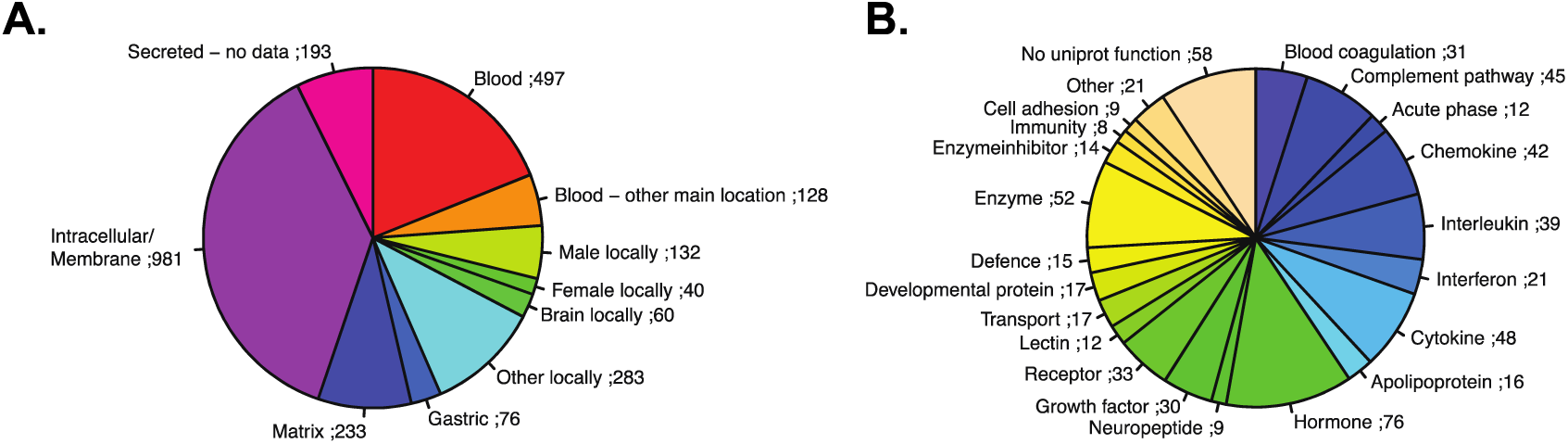
Annotation and analysis of the human secretome. **A.** The annotation of the 2,623 proteins defined as the human secretome based on bioinformatics criteria. The proteins are classified according to their predicted final localization. If several isoforms exist with different predicted localization, only one location is annotated per gene. **B.** The predicted function of the 625 proteins classified as blood proteins.

We subsequently analyzed the 625 predicted blood proteins from the groups “Blood” and “Blood - other main location”. **Fig. 1B** shows functional classification of these proteins, and the major groups consist of cytokines, interleukins, interferons and chemokines (n=150), complement and coagulation factors (n=76), hormones (n=76), growth factors (n=30), and enzymes (n=52). 58 proteins had no function keyword in UniProt and these are interesting targets for in-depth studies to investigate their possible role in blood. An analysis of the tissue-specificity based on the Human Protein Atlas transcriptomics data showed that a majority of the blood proteins show a tissue-restricted expression. Interestingly, the fraction of “housekeeping” proteins expressed in all tissues is only 22%, which is much lower than the fraction for all protein-coding genes (38%)^1^. Further analysis of the tissue-specificity of the transcriptome of the tissue-enriched blood proteins reveals that a large majority of them are expressed in the liver. This is not unexpected since the liver is known for production of many of the plasma proteins^7^.

### CHO cell factory for the production of the human secretome

We decided to set up a standardized expression system based on the mammalian CHO cell host system combined with semi-stable transfection of clones generated by synthetic biology. To enable production and purification of all secreted proteins, genes encoding the target proteins were synthesized (**Fig. 2A**) and all proteins were transiently produced in mammalian CHO cells using the QMCF Technology^8^ and subsequently purified using an antibody-based chromatography resin with calcium ion-dependent affinity for the HPC4 tag, included in the C-terminus of the recombinant protein. This tag enables mild elution by the use of a chelating elution buffer. The purity and protein identity of the various target proteins were analyzed by SDS-PAGE, Western blot and MS/MS (**Fig. 2B**). To simultaneously determine concentration and identity of the produced proteins, an MS-based method for absolute quantification was developed based on spike-in of stable isotope-labeled protein fragments (QPrESTs)^9, 10^ corresponding to the target proteins. A comparison between the targeted proteomics analysis and the standard operating procedure to determine protein concentrations using spectrophotometric principles demonstrated that the latter method often gives misleading results of both over- and under-estimation of the concentration of the target protein. We have therefore determined the absolute concentration of each protein using the MS based workflow.

**Figure 2.**
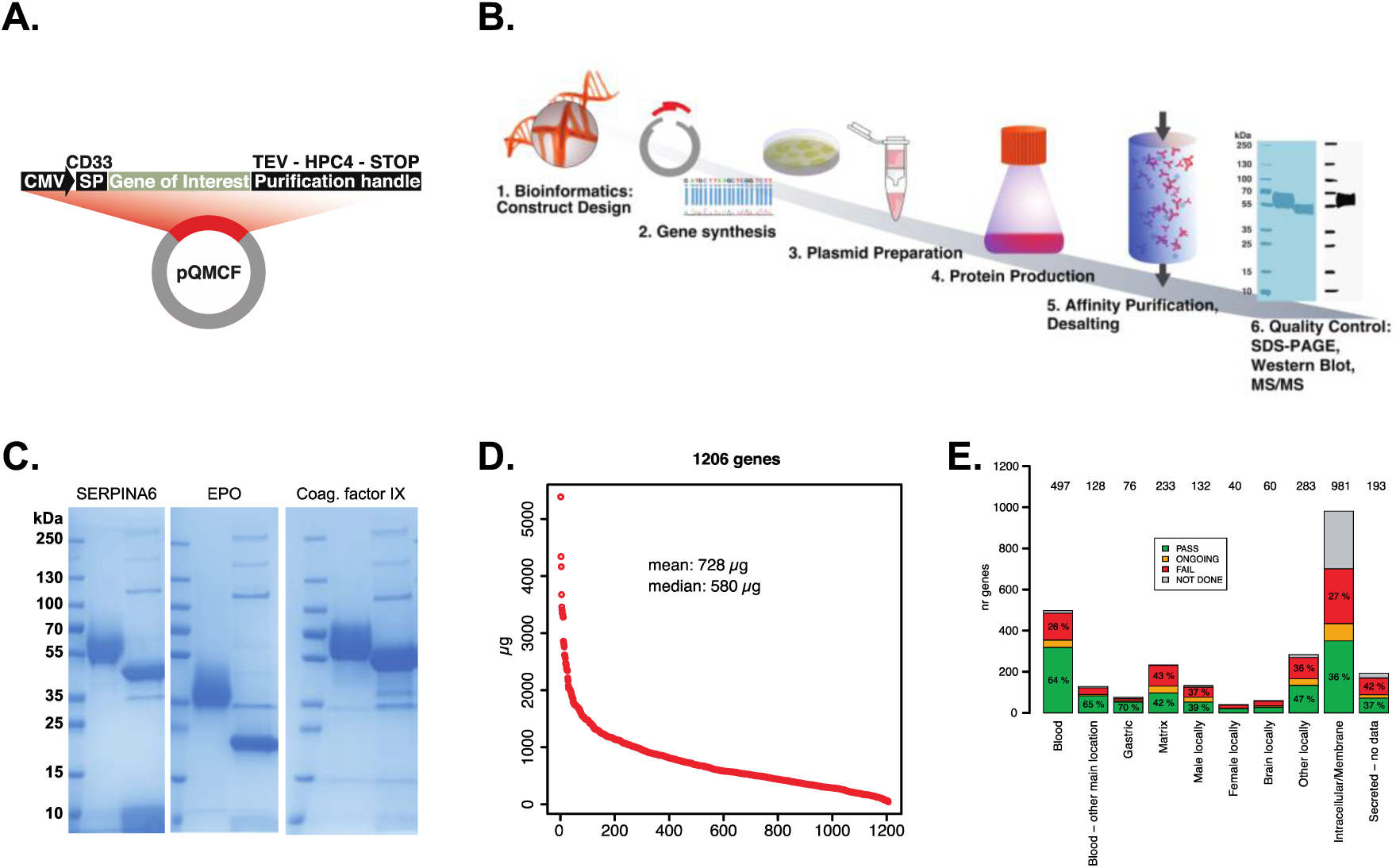
Bioproduction of the human secretome in CHO cells. **A.** All proteins in the HSP are produced with an N-terminal signal peptide, CD33, and a C-terminal purification handle based on the HPC4 purification tag^11^. Between the purification tag and the protein, a DNA sequence encoding a TEV protease site^12^ was inserted to allow for cleavage of the tag, if needed. The CMV promoter is used to control the protein production. **B.** The first step in the standardized high-throughput protein production pipeline used in the HSP is construct design. The constructs are then synthesized and cloned into the expression plasmid. All plasmids are prepared and sequence verified before protein production. The HPC4-tagged target proteins are then purified using an automated affinity purification setup. Purified proteins are identified and quantified with MS/MS and purity and glycosylation pattern are determined using SDS-PAGE and Western Blot. **C.** A SDS-PAGE showing three representative proteins after purification. The first lane shows the purified protein sample and second lane the protein after deglycosylation. Arrows indicate the enzymes from the deglycosylation kit. **D.** The amount of the different target proteins generated by the stream-lined approach. **E.** The success rates for the CHO clones are shown. The overall success rate is approximately 50%, although differing considerably between different groups based on predicted localization.

Many of the secreted proteins are post-translationally modified during translocation through the secretory pathway. Therefore, the degree of glycosylation was assessed using a deglycosylation enzyme kit. This is exemplified in **Fig. 2C** showing a heterogenous size band after CHO production and a single band of expected size after deglycosylation. This shows that the CHO host system often yields proteins with differential glycosylation patterns, which is not unexpected and a phenomenon that has been reported before^13^.

Based on this standardized pipeline, clones corresponding to 2,265 genes were transfected to CHO host cells and the production of corresponding full-length target protein was determined. The amounts of proteins produced were highly variable (**Fig. 2D**) and for almost half of the proteins, no or little target protein was recovered. The success rate varied considerably between the different protein classes defined above (**Fig. 2E**) with the highest success rates for gastric and blood proteins (70% and 64%, respectively), while proteins expected to be intracellular or membrane-bound show the lowest success rate (36 %).

### Variation in achieved protein amount cannot be explained by mRNA abundance

Among the library of CHO cells producing the human secretome, 96 representative CHO cell lines were subjected to transcriptomics analysis. We visualized the global transcriptome usage of the CHO cells using a Proteomap^14^ (**Fig. 3A**), wherein mRNA abundance is summarized into blocks of cellular processes with the area representing the fraction of the transcriptome dedicated to a particular cellular process. On average, the transgenes are among the most highly expressed genes in the CHO cell transcriptome, taking up ∼3% of the transcriptome and the levels of mRNA were comparable among cell lines, regardless of whether the cell lines secreted high, low, or no amounts of recombinant protein (**Fig. 3B**). Additionally, the major cellular processes remain relatively stable across different tiers of producers and interestingly, the expression level of genes associated with translation show slightly higher expression in cell lines with the lowest recombinant protein yield. Incidentally, cellular processes associated with vesicular transport were down-regulated in non-producers.

**Fig. 3.**
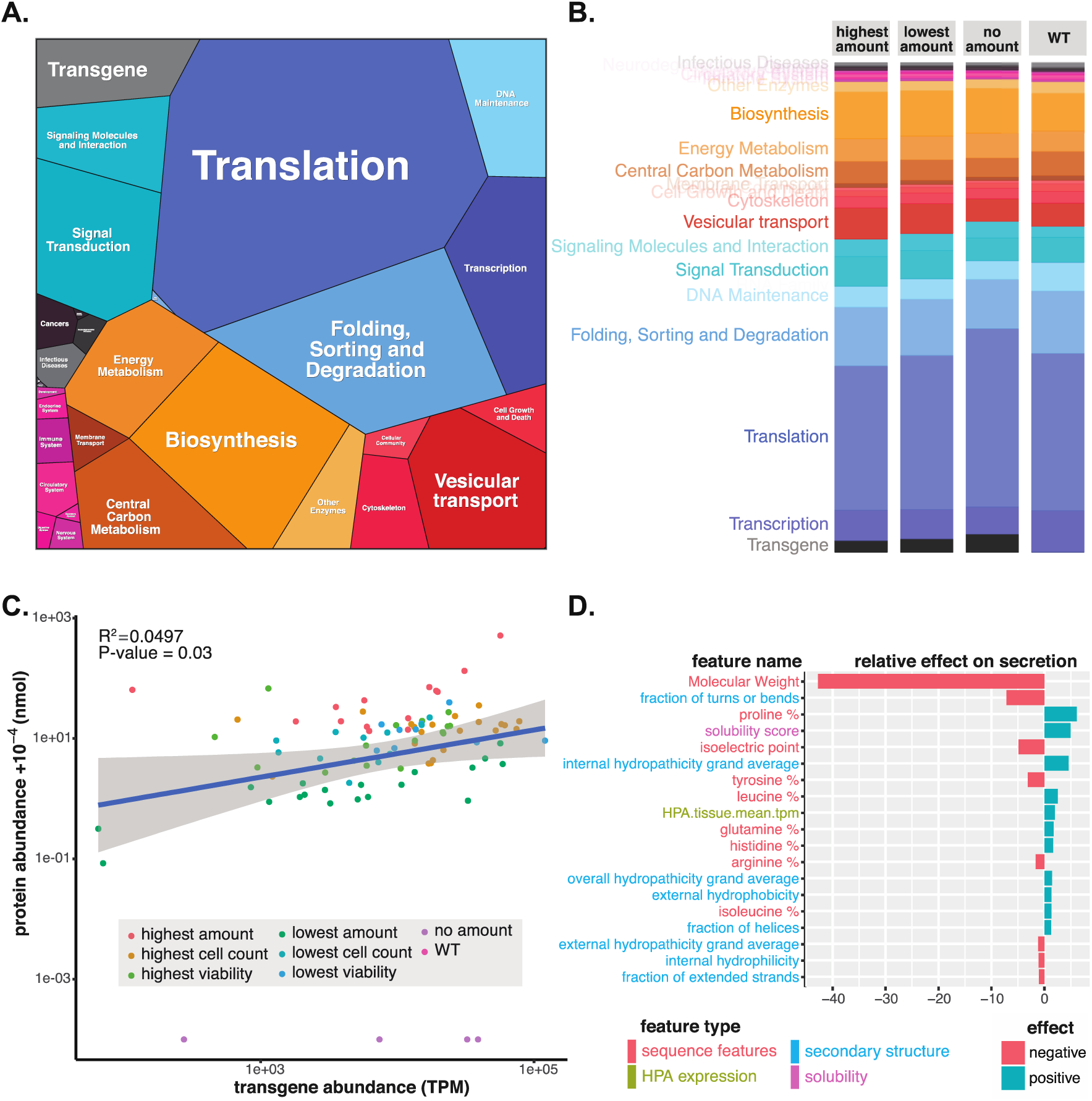
The contribution of transgene mRNA level, host cell transcriptome, and protein structural properties on the yield of recombinant proteins in CHO cells. **CHO cells expressing the entire human secretome were developed, and RNA-Seq was conducted on 96 representative clones, each secreting a different human protein. A.** The cells expressed a wide range of transcripts supporting a wide range of cellular processes. The cells dedicated a substantial amount of resources on mRNAs associated with translation and protein synthesis (blue). **B.** Overall, the transcriptome usage is mostly consistent across cell lines showing different levels of recombinant protein production. **C.** The transgene abundance (TPM) of each recombinant gene was compared to the secreted protein abundance (nmol). **D.** To quantify the impact of secreted protein characteristics and their effects on secretion, a machine learning pipeline was used to identify features of the secreted proteins that contribute the most (both positively and negatively) to protein secretion.

We further evaluated if the variation in protein production yields can be explained by transgene mRNA levels. The transgene mRNA levels only explain 5% (**Fig. 3C**) of the variance in recombinant protein productivity, compared to previous reports of ∼40% for endogenous genes in mammalian cells^15, 16^. In the 96 samples studied, the four cell lines that did not secrete detectable levels of recombinant protein have moderate to high levels of transgene mRNA. This suggests that difficult-to-express proteins are less limited by transcript level, compared to other factors.

We further examined how other transcriptomic determinants in the host cells impacted the amount of protein achieved. To evaluate if any particular pathway differentiated between the cell lines that gave high vs. low amount of recombinant protein product, we conducted Gene Set Enrichment Analysis (GSEA)^17, 18^ and found several dysregulated cellular responses. This includes genes targeted by KRAS activation, whose expression was significantly down-regulated in high producers. As KRAS contributes to the Warburg effect^19^, the reduction in KRAS signaling in high producers may improve cell growth, contributing to higher recombinant protein yields.

### Relating a protein’s physical properties to its characteristic expression

We further evaluated if attributes of each secreted protein could explain protein titers. Thus, to determine secretome-specific determinants of protein secretion, we used machine learning to model how the features of the proteins affect the protein yield (**Fig. 3D**). Note that the transgene mRNA level was excluded from this analysis to focus on structural features of the recombinant proteins. Among more than 150 features curated for the secreted proteins, the molecular weight of the proteins impacts achieved protein amounts the most. Large proteins pose additional demands, as they have more post-translational modifications, provide more opportunities for misfolding and aggregation, more difficulty of trafficking, and impose increased energy consumption in their production. The amount protein attained is also impacted considerably by other features, such as the fraction of the protein sequence to be in structural turns or bends, the amount of proline, hydrophobicity, etc. Thus, these attributes provide features to consider when producing recombinant proteins.

### Rescue of difficult proteins in human cell line HEK293

We subsequently decided to investigate if proteins failed in the CHO cell factory could be rescued by expression in a human cell line. Constructs that previously failed in the CHO cell production were transfected to HEK293 and a standard cultivation procedure was employed (**Fig. 4A**). Many proteins showing considerable degradation in CHO were successfully produced in the human cells with some examples shown in **Fig. 4B** and **4C**. Out of 129 protein constructs that earlier failed in the CHO cell factory due to degradation, 88 (68%) were successfully rescued by changing the production host from CHO to HEK293 (**Fig. 4D**). The results suggest that using a human cell line would be an attractive option for proteins difficult to express in the standard biomanufacturing CHO host cell line.

**Figure 4.**
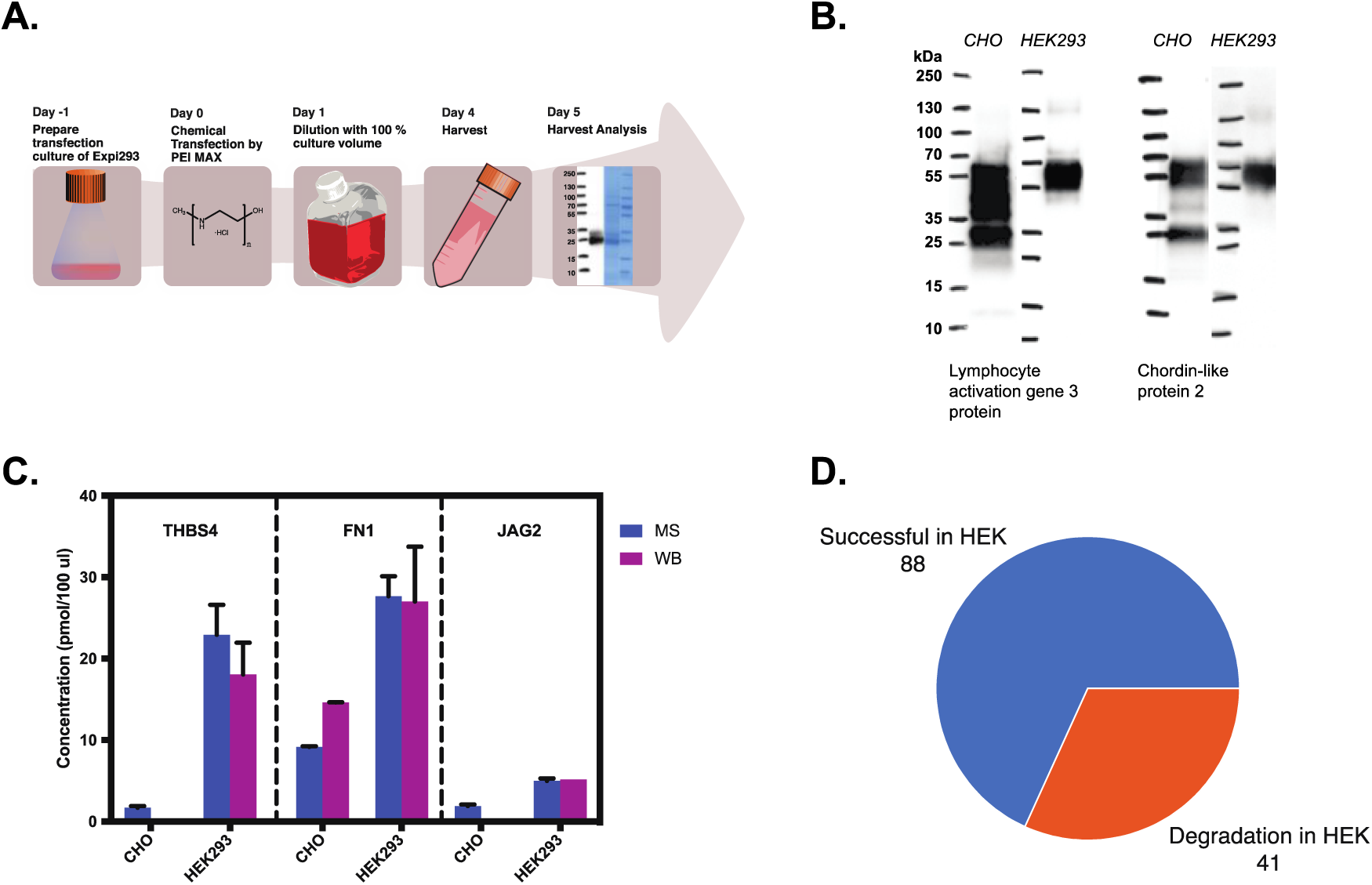
Rescue of difficult proteins in human cell line HEK293. **A.** Currently used protein production protocol to rescue proteins that due to degradation failed in the standard pipeline. This protocol was chosen after evaluation of different versions of HEK293 cells and different protocols. **B.** Examples of Western Blots with samples from the harvest in CHO (left WB) and HEK293 (right WB). **C.** Absolute concentrations measured by MS and WB of three different proteins produced in CHO and HEK293. **D**. The overall results for the 129 proteins that failed due to protein degradation in the standard CHO workflow and were subsequently expressed using the rescue protocol in the human cell line HEK293.

### Use of the secretome resource for phenotypic assays

A set of the produced proteins were further used to explore the secretome resource for stimuli-induced changes in phenotypic screens using physiologically relevant cells to aid in the development of drug candidates as described before^20-23^. The “in-house” generated library of proteins was used to explore the prevention of loss of β-cell function in the pancreas (**Fig. 5A**). By screening proteins from 763 different secretome genes, we found that several fibroblast growth factors triggered human β-cell dedifferentiation, including the positive control FGF2 (**Fig.5B**). Interestingly, FGF9 shows a higher activity than the control, but also FGF1, FGF4 and FGF18 show activity above background. To our knowledge, the activity of several of these fibroblast growth factors have not been described before in this context. Single cell expression analysis of FGFR expression on primary human islets shows that FGFR1 is dominantly expressed (**Fig 5C**). Taken together, the data suggests that therapeutic intervention could be achieved by inhibition of FGFR1 signaling with no additional signaling pathways identified that maintain the dedifferentiated state of β cells.

**Figure 5.**
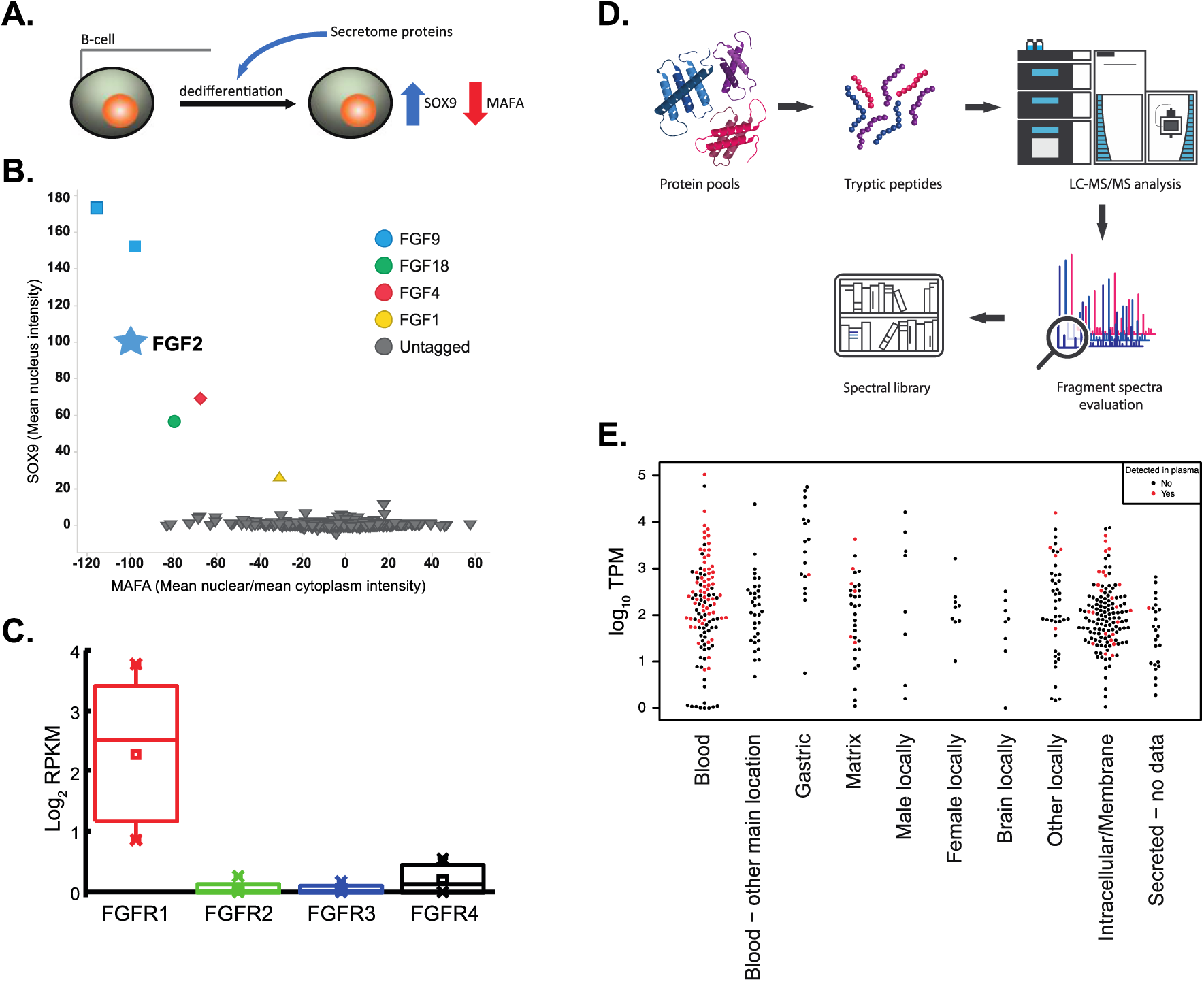
Using the secretome resource for developing phenotypic and proteomics assays. A. Schematic view of a β-cell dedifferentiation assay. The library was screened in a human β-cell line EndoC-βH1 to identify secreted proteins that affect the differentiation state of the cells^24^. Transcription factors SOX9 and MAFA were used as markers for the dedifferentiated and differentiated state, respectively. EndoC-βH1 cells were treated with secretome proteins or positive control (FGF2) at three different concentrations. B. Response to FGF2 on both MAFA and SOX9 was set as 100% (indicated by star). Retesting of initial proteins that downregulated MAFA and upregulated SOX9 in 10-point concentration responses confirmed FGF9, FGF4, FGF18 and FGF1 as inducers of EndoC-βH1 dedifferentiation with FGF9 being more active than the FGF2 positive control. C. RNAseq analysis of FGFR expression in human primary β-cells isolated from human pancreatic islets. Human islet samples (85%–95% pure) were dissociated and distributed by FACS into 384-well plates before expression analysis was performed. D. Schematic view of the proteomics assay development. Proteins were combined into pools of eight and digested with trypsin before analysis in shotgun-MS mode. Peaks of detected peptides were manually inspected and their fragmentation spectra were used for DIA data extraction. E. Beeswarm plot showing the secreted proteins for which DIA assays could be established and whether they could be detected (red) or not (black) in human plasma analyzed through DIA. The proteins are divided in to their respective secretome classification categories and the y-axis is showing the mRNA expression level as the maximum TPM value based on 37 normal human tissues.

### Use of the secretome resource to develop proteomics assays

Data-independent acquisition (DIA) mass spectrometry can provide highly quantitative MS2-data for thousands of proteins in a single analysis, but requires libraries of peptide fragmentation spectra and retention times for identification of peptide peaks during the data analysis step^25, 26^. These fragmentation spectra are often based on shotgun proteomics analysis of complex samples which might result in low quality spectra where low abundant proteins are often missing. Therefore, it was decided to make use of the secretome resource to develop a high-quality spectral library by analyzing trypsin digested equimolar pools of the purified proteins in shotgun mode (**Fig. 5D**). In total, secreted proteins corresponding to 474 unique genes were screened and the fragmentation spectra were manually verified using the Skyline software^27^. For 434 of the proteins, DIA assays could be created based on at least one tryptic peptide, while 40 of the proteins did not result in any good peptide fragment spectra. We then analyzed human plasma samples, and peptide data corresponding to 96 proteins could be extracted using the developed spectral library, while 338 proteins could not be detected using this DIA proteomics analysis (**Fig. 5E**). As expected, proteins that are actively secreted into the blood were detected in plasma to a larger extent than proteins in any other category, although some proteins belonging to the non-blood categories, especially proteins with relatively high mRNA levels in the highest expressed tissue, were detected in plasma. It is noteworthy that although many of the gastric proteins have very high expression in at least one tissue only one the proteins from this category could be detected in blood, supporting a notion that gastric proteins are not secreted to the blood.

## Discussion

The human secretome is a highly interesting group of proteins both for studies of human biology and as targets for the development of new drugs and diagnostics. Here, we report an annotated list of the human secretome that includes a prediction of the final location of each of these proteins. The annotation has been based on bioinformatics analysis and available literature and provides for the first time a comprehensive view of the actively secreted proteins in humans. Out of the 2,623 proteins defined as the secretome, almost a thousand are suggested to be intracellular and almost 800 are transported to local compartments as either extracellular matrix or specialized tissues, such as the male/female organs. Another 80 proteins are secreted to the gastrointestinal tract, thus leaving only about 600 proteins predicted to be actively secreted to human blood. This list will be revised when more information will be available in the future, but it is noteworthy that the number of predicted blood proteins are surprisingly low, corresponding to only 3% of all human protein-coding genes. It is however reassuring that the list of blood proteins contains the expected classes of plasma proteins, such as cytokines, apolipoproteins, coagulation and complement factors, hormones, enzymes and growth factors.

The human blood proteome comprises proteins that are either actively secreted from various human tissues (the secretome), but also proteins that are leaked by the millions of cells undergoing cell death at any given moment. However, the actively secreted proteins constitute the vast majority of these blood proteins and as few as ten of them make up more than 90% of the total protein mass in human blood. The dynamic range of the human secretome in the blood covers more than 10 orders of magnitude, from albumin at 45 mg per mL blood to low abundant proteins, such as the cytokine IL-1, at less than 5 pg per mL. A quest for the future is thus to provide comprehensive assays allowing the analysis of all these proteins in a multiplex fashion despite their huge dynamic range.

Here, a high-throughput mammalian cell factory was set-up with recombinant expression in CHO cells of full-length gene versions of each of the potentially secreted proteins. More than 60% of the genes coding for blood proteins were successfully recovered from the culture medium, while the success rates for the other categories of secreted proteins were somewhat lower giving a total one-pass success rate of approximately 50%. An omics analysis of the CHO cells demonstrated that transcriptional load was not a bottle-neck for the production, while biochemical features such as molecular weight and molecular bends of the target protein were found to contribute significantly to the success rate of the bioproduction. For proteins showing degradation in CHO cells, a large majority could be rescued in the human cell line HEK293, demonstrating that this is an attractive alternative to CHO production for biomanufacturing of human proteins. More than 1,200 human secreted proteins were produced and successfully purified. In conclusion, the human secretome has been mapped and characterized to facilitate further exploration of the human secretome.

## Acknowledgments

Main funding was provided from the Novo Nordisk Foundation, the Knut and Alice Wallenberg Foundation, AstraZeneca and NIH grant NIGMS (R35 GM119850). We acknowledge the entire staff of the Human Protein Atlas program and the Science for Life Laboratory for their valuable contributions.

## References

1. Uhlen, M. et al. Proteomics. Tissue-based map of the human proteome. Science 347, 1260419 (2015).

2. Altay, G. & Peters, B. Gene Regulatory Cross Networks: Inferring Gene Level Cell-to-Cell Communications of Immune Cells. bioRxiv (2018).

3. Pelham, H.R. The retention signal for soluble proteins of the endoplasmic reticulum. Trends in biochemical sciences 15, 483–486 (1990).

4. Gould, S.J., Keller, G.A., Hosken, N., Wilkinson, J. & Subramani, S. A conserved tripeptide sorts proteins to peroxisomes. The Journal of cell biology 108, 1657– 1664 (1989).

5. Legakis, J.E. & Terlecky, S.R. PTS2 protein import into mammalian peroxisomes. Traffic 2, 252–260 (2001).

6. Schatz, G. The protein import system of mitochondria. The Journal of biological chemistry 271, 31763–31766 (1996).

7. Kampf, C. et al. The human liver-specific proteome defined by transcriptomics and antibody-based profiling. FASEB journal: official publication of the Federation of American Societies for Experimental Biology 28, 2901–2914 (2014).

8. Silla, T. et al. Episomal maintenance of plasmids with hybrid origins in mouse cells. Journal of virology 79, 15277–15288 (2005).

9. Zeiler, M., Straube, W.L., Lundberg, E., Uhlen, M. & Mann, M. A Protein Epitope Signature Tag (PrEST) library allows SILAC-based absolute quantification and multiplexed determination of protein copy numbers in cell lines. Molecular & cellular proteomics: MCP 11, O111 009613 (2012).

10. Edfors, F. et al. Immunoproteomics using polyclonal antibodies and stable isotope-labeled affinity-purified recombinant proteins. Molecular & cellular proteomics: MCP 13, 1611–1624 (2014).

11. Stearns, D.J., Kurosawa, S., Sims, P.J., Esmon, N.L. & Esmon, C.T. The interaction of a Ca2+-dependent monoclonal antibody with the protein C activation peptide region. Evidence for obligatory Ca2+ binding to both antigen and antibody. The Journal of biological chemistry 263, 826–832 (1988).

12. Dougherty, W.G., Cary, S.M. & Parks, T.D. Molecular genetic analysis of a plant virus polyprotein cleavage site: a model. Virology 171, 356–364 (1989).

13. Croset, A. et al. Differences in the glycosylation of recombinant proteins expressed in HEK and CHO cells. Journal of biotechnology 161, 336–348 (2012).

14. Liebermeister, W. et al. Visual account of protein investment in cellular functions. Proceedings of the National Academy of Sciences of the United States of America 111, 8488–8493 (2014).

15. Wilhelm, M. et al. Mass-spectrometry-based draft of the human proteome. Nature 509, 582–587 (2014).

16. Schwanhausser, B. et al. Global quantification of mammalian gene expression control. Nature 473, 337–342 (2011).

17. Subramanian, A. et al. Gene set enrichment analysis: a knowledge-based approach for interpreting genome-wide expression profiles. Proceedings of the National Academy of Sciences of the United States of America 102, 15545–15550 (2005).

18. Mootha, V.K. et al. PGC-1alpha-responsive genes involved in oxidative phosphorylation are coordinately downregulated in human diabetes. Nature genetics 34, 267–273 (2003).

19. Weinberg, F. et al. Mitochondrial metabolism and ROS generation are essential for Kras-mediated tumorigenicity. Proceedings of the National Academy of Sciences of the United States of America 107, 8788–8793 (2010).

20. Lin, H. et al. Discovery of a cytokine and its receptor by functional screening of the extracellular proteome. Science 320, 807–811 (2008).

21. Swinney, D.C. & Anthony, J. How were new medicines discovered? Nature reviews. Drug discovery 10, 507–519 (2011).

22. Eder, J., Sedrani, R. & Wiesmann, C. The discovery of first-in-class drugs: origins and evolution. Nature reviews. Drug discovery 13, 577–587 (2014).

23. Gonzalez, R. et al. Screening the mammalian extracellular proteome for regulators of embryonic human stem cell pluripotency. Proceedings of the National Academy of Sciences of the United States of America 107, 3552–3557 (2010).

24. Diedisheim, M. et al. Modeling human pancreatic beta cell dedifferentiation. Molecular metabolism 10, 74–86 (2018).

25. Venable, J.D., Dong, M.Q., Wohlschlegel, J., Dillin, A. & Yates, J.R. Automated approach for quantitative analysis of complex peptide mixtures from tandem mass spectra. Nature methods 1, 39–45 (2004).

26. Gillet, L.C. et al. Targeted data extraction of the MS/MS spectra generated by data-independent acquisition: a new concept for consistent and accurate proteome analysis. Molecular & cellular proteomics: MCP 11, O111 016717 (2012).

27. MacLean, B. et al. Skyline: an open source document editor for creating and analyzing targeted proteomics experiments. Bioinformatics 26, 966–968 (2010).

